# Parallel sequencing lives, or what makes large sequencing projects successful

**DOI:** 10.1101/136358

**Authors:** Javier Quilez, Enrique Vidal, François Le Dily, François Serra, Yasmina Cuartero, Ralph Stadhouders, Thomas Graf, Marc A. Marti-Renom, Miguel Beato, Guillaume Filion

**Author notes:** Email addresses: JQ.

## Abstract

T47D_rep2 and b1913e6c1_51720e9cf were two Hi-C samples. They were born and processed at the same time, yet their fates were very different. The life of b1913e6c1_51720e9cf was simple and fruitful, while that of T47D_rep2 was full of accidents and sorrow. At the heart of these differences lies the fact that b1913e6c1_51720e9cf was born under a lab culture of Documentation, Automation, Traceability, Autonomy and compliance with the FAIR Principles. Their lives are a lesson for those who wish to embark on the journey of managing high throughput sequencing data.

## The beginning

Linda worked hard to produce a Hi-C sample in T47D cells. Upon submitting the sample for sequencing, she remembered the motto of the lab: “Make DATA more FAIR”. The team had established lab-wide habits of Documentation, Automation, Traceability and Autonomy of experimenters. The old-timers insisted that human interfaces are always the weak link. “Every time a project fails, someone is typing on a keyboard… or does not bother to”. The metadata must be accurate, the code must be readable, the data must be tidy. Technology helps, but this is mostly a matter of attitude. Not only had such attitude improved the performance of the lab but it also paved the way to meet international quality standards as those defined by the FAIR Principles [1].

Linda filled in the metadata on a low-key online Google Form. The lab had chosen this option among many others because experimenters found it the easiest. Filling the form was quick: they had to click on items from drop-down lists. As she pressed “Submit”, a shared Google Sheet was immediately updated and she received the name b1913e6c1_51720e9cf that uniquely identified her sample. These unnatural names had first left her skeptical, but she could now see the benefits of that system to collect the metadata and trace sequencing samples. She remembered the meetings with the bioinformaticians in an attempt to make the data more FAIR [1]. “A project is as good as its metadata; you will see the benefit only after a year or two” they kept telling.

Meanwhile in another lab, Pedro also worked hard to produce a Hi-C sample in T47D cells. Things had gone wrong in the past, but this time all the quality controls looked good. He proudly wrote “T47D_rep2” on the tube and gave it to the sequencing facility. All the information *he* considered relevant was in his notebook.

By a strange coincidence, both Linda and Pedro soon found a new position. They left their respective institutes without finishing their project.

## Life after turn-over

Simon was the bioinformatician in charge of analyzing T47D_rep2. He was not happy that Pedro left the institute, because he had questions about the sample. As he meant to save the files in the shared repository, he realized that there were already four samples called “T47D_rep2” in different directories. Simon facepalmed and headed for the wet lab. Fortunately, Janet knew something about it: “Some of these are my experiments; the others are Pedro’s. Despite the modest sequencing coverage, he found interesting changes in the genome structure when treating with hormone, so he repeated the experiments to obtain higher coverage”. Looking into Pedro’s notes, Simon saw that indeed the sequencing quality of the raw reads was very poor, hence the newest sample “T47D_rep2”. At long last, Simon had an idea of what “T47D_rep2” was…

Meanwhile, Paul, the bioinformatician in charge of analyzing b1913e6c1_51720e9cf pulled the record from the database where the metadata in the Google Sheet were automatically dumped. The online spreadsheet was a convenient frontend for the experimenters, but the database offered a more programmatic access to the metadata — plus it was an additional backup layer. On his end, Paul launched the mapping pipeline and performed several downstream analyses that Chloe requested. He documented the procedure in the Jupyter electronic notebook he created for the analysis. The production code was run in Docker containers and pushed to a GitHub repository. The notebooks helped him (or anyone else) keep track of the analyses in a readable format, while Docker virtual machines allowed him (or anyone else) to run the code on different machines without the hassle of installing countless libraries. Finally, GitHub was as much a backup as a way to share his work.

Chloe examined the results in the online report she received from Paul and performed some additional analyses with an R Shiny web application to inspect the Hi-C data processed in the lab. It had taken some time to implement it, but now the benefits were clear: Paul could focus on other things than running basic analyses for all the lab members and, meanwhile they were more autonomous. This last analysis provided further evidence supporting their hypothesis, so Chloe was ready to polish their manuscript. Each analysis performed by Paul was allocated in a directory with a traceable name, a clear content structure and permanently accessible in the FTP site of the lab. Therefore, Chloe knew where to find the figures and tables that she needed, updated the Methods section with the information written in the report and she was even able to provide the scripts and parameter values used in the analysis as a GitHub repository — she knew that editors were getting more and more serious about reproducibility.

## The reviews

Chloe was very happy to hear their manuscript received positive comments from the reviewers. The only obstacle to publication seemed to be Reviewer #3, who asked to replicate the findings in an independent larger dataset that had been recently published. Tough but fair. Chloe panicked about having to analyze almost 100 samples in so little time; during the project they had generated a smaller number of samples and analyzed them over time, so she worried that it would take too long. Paul reassured her: all she had to do was prepare the metadata for the new dataset, as Linda had done for b1913e6c1_51720e9cf. Then, a simple command would execute the pipeline for the ∼100 samples as effortlessly as for a single one, and all the required information would be retrieved automatically from the database of metadata. Running the pipeline could be parallelized in the multiple cores available in the computing cluster of the institute, so all samples were processed within a few days. In the meantime, he would start preparing the submission of the data to a public repository: a simple search within the structured directories allocated for the FASTQ and the contact matrix files as well as a selection of entries from the database of metadata would do much of the work. Lastly, Paul checked that the manuscript complied with the FAIR Principles [1]. Findability and accessibility: the data and metadata were linked by the unique sample identifier and uploaded to GEO, the code was pushed to GitHub and the URL to both repositories available in the manuscript. Interoperability: the Docker containers used to run the pipelines were pushed to Docker Hub. Reusability: the metadata was complete and the data procedures were well documented.

Meanwhile, Simon was far from publication. Overall, the preliminary results of Pedro were not confirmed in the new high-coverage samples. Simon scavenged the directories looking for the code used to generate the plots he had seen, those that indicated a clear effect of hormone treatment on the genome structure. Unfortunately, the workflow of the analysis and the specific parameter values were not documented. Perhaps his predecessors had forgotten to remove PCR duplicates? And how did they correct for multiple testing, if at all? After guessing where to find the older raw data, Simon processed the initial dataset with his analysis pipeline but the differences between the old and new datasets remained. Simon facepalmed. He knew too well that trouble was only starting…

## Behind the scene

The human factor is the greatest hurdle to reaching the standard of the FAIR Principles [1]. People change their mind, they resist change, they follow their own rules and they plan for the short term. As an insurance against fiasco (**Table 1**), a scientific team must develop habits and tools for sharing data and analyses. The main idea is to limit or control human intervention by automating every step.

**Table 1.** Challenges associated to the accelerated accumulation of high throughput sequencing data. As storified with the lives of b1913e6c1_51720e9cf and T47D_rep2, managing and analyzing the growing amount of sequencing data presents several challenges.

| Challenge | Impact | Consideration |
| --- | --- | --- |
| Mislabelled raw sequencing data | • Underpowered analysis<br>• Erroneous results<br>• Loss of data, time and resources | Check unassigned reads and sequencing index concordance |
| Poor sample description | • Prevents data processing and quality control<br>• Incorrect analysis and results<br>• Lack of reproducibility<br>• Delays publication | Metadata collection |
| Unsystematic sample naming | • Duplicated or similar names<br>• Ambiguous identification<br>• Precludes computational treatment<br>• Data disclosure | Sample identifier scheme |
| Untidy data organisation | • Data cannot be found<br>• Time consumption<br>• Inability to automate searches | Structured and hierarchical data organisation |
| Yet another analysis | • Repeated manual execution of analyses<br>• Incapability to deconvolute analysis producing different results<br>• Compulsory linear execution | Scalability, parallelization, automatic configuration and modularity |
| Undocumented procedures | • Poor understanding of results<br>• Irreproducibility<br>• Hampers catching errors | Documentation |
| Data overflow | • No access to data<br>• Size and number of files make individual inspection inefficient | Interactive web applications |

1. The absolute priority is metadata collection. We propose a scheme for collection and file naming (**Figure 1a** and **Additional file 1**), but any system will do, as long as it is (i) agreed upon and understood by people using it, (ii) backed up automatically, (iii) future-proof and (iv) there is someone responsible for maintenance and validation of the metadata.
2. The second priority is to locate the data and the analyses. We propose a hierarchical organization that can evolve according to future needs (**Figure 1b**). Again, any scheme with the properties above will do.
3. Next, the analyses must be documented. Here a flurry of tools help the analysts keep track of and organize their work as it unfolds. The most popular are Jupyter for Python and Rstudio for R. Here we recommend using widely accepted tool kits as this facilitates sharing between the members of the team and the rest of the world.
4. Such tools partly address the next priority, which is reproducibility. However, today we can go one step further with virtual machines. In this area, Docker has taken the lead and we recommend developing ground up production scripts and exploratory analyses in Docker containers.
5. Finally, experimenters should be empowered to perform basic analyses. The most efficient teams are made of specialists, so researchers should do what they are expert at (or become expert at what they do). But bioinformatics is fast becoming “common knowledge”. Building interfaces for standard analyses is a way to free bioinformaticians to focus on the most technical parts of the project, while allowing all the members to contribute to the analyses. Many modern tools such as R Shiny can help build such interfaces. Here, the most important is that the developer be proficient with the chosen tool, and that they users understand how to use the interface.

**Figure 1.**
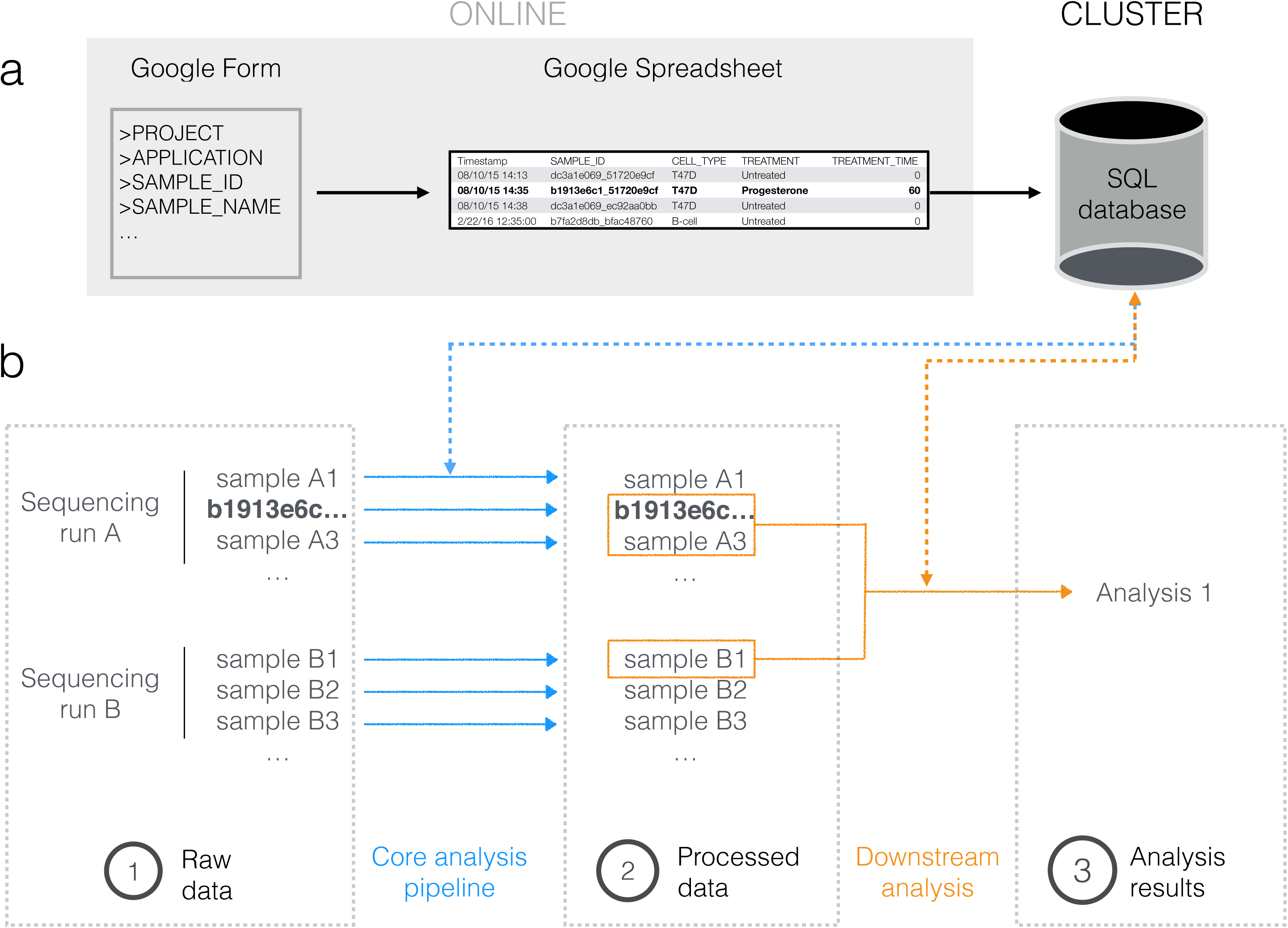
A traceable life for b1913e6c1_51720e9cf. **(a)** The metadata for b1913e6c1_51720e9cf were collected via an online Google Form and stored both online (Google Sheet) and in a local SQL database. A good metadata collection system should be (i) short and easy to complete, (ii) instantly accessible by authorized users and (iii) easy to parse for humans and computers. **(b)** b1913e6c1_51720e9cf was sequenced along with other samples, whose raw sequencing data were located in a directory named after the date of the sequencing run. There one could find the FASTQ files containing the sequencing reads from b1913e6c1_51720e9cf as well as information about their quality; no modified, subsetted or merged FASTQ file was stored to ensure that analyses started off from the very same set of reads. In a first step, the raw data of b1913e6c1_51720e9cf were processed with the Hi-C analysis pipeline, which created a “b1913e6c1_51720e9cf” directory at the same level where all processed Hi-C samples were located. “b1913e6c1_51720e9cf” had multiple subdirectories that stored the files generated in each of the steps of the pipeline, the logs of the programs and the integrity verifications of key files. Moreover, such subdirectories accounted for variations in the analysis pipelines (e.g. genome assembly version, aligner) so that data were not overwritten. In a second step, processed data from b1913e6c1_51720e9cf and other samples were used to perform the downstream analyses Chloe asked Paul. Within the directory he allocated to her analyses, Paul created a new one called “2017-03-08_hic_validation” with the description of the analysis along with the scripts used and the tables and figures generated.

Data accumulates at a rapid pace in life sciences (**Additional file 2**), and stories similar to that of b1913e6c1_51720e9cf and T47D_rep2 have taken place in many research groups (**Additional files 3-5**). We propose that data-producing teams focus on Documentation, Automation, Traceability and Autonomy as main priorities, with the purpose of being “human-proof”. The scheme implemented in our own projects is shown in **Figures 1-2**, and the tools are listed in **Table 2**. To illustrate our recommendations, we also provide a didactic data set (the actual sample b1913e6c1_51720e9cf) at the following link: https://github.com/4DGenome/parallel_sequencing_lives.

**Table 2.** Tools used in the story.

| Tool | Usage | Website |
| --- | --- | --- |
| Docker | Interoperability | https://www.docker.com/ |
| Docker Hub | Repository for Docker containers | https://hub.docker.com/ |
| GEO | Repository for high-throughput genomics data | https://www.ncbi.nlm.nih.gov/geo/ |
| GitHub | Version control and backup of code | https://github.com/ |
| Google Forms and Sheets | Online collection and display of metadata | https://www.google.com/forms/about/ |
| Jupyter Notebook | Document procedures and perform analysis | http://jupyter.org/ |
| R Shiny | Deploy web applications | https://shiny.rstudio.com/ |
| R Studio | Document procedures and perform analysis | https://www.rstudio.com/ |

**Figure 2.**
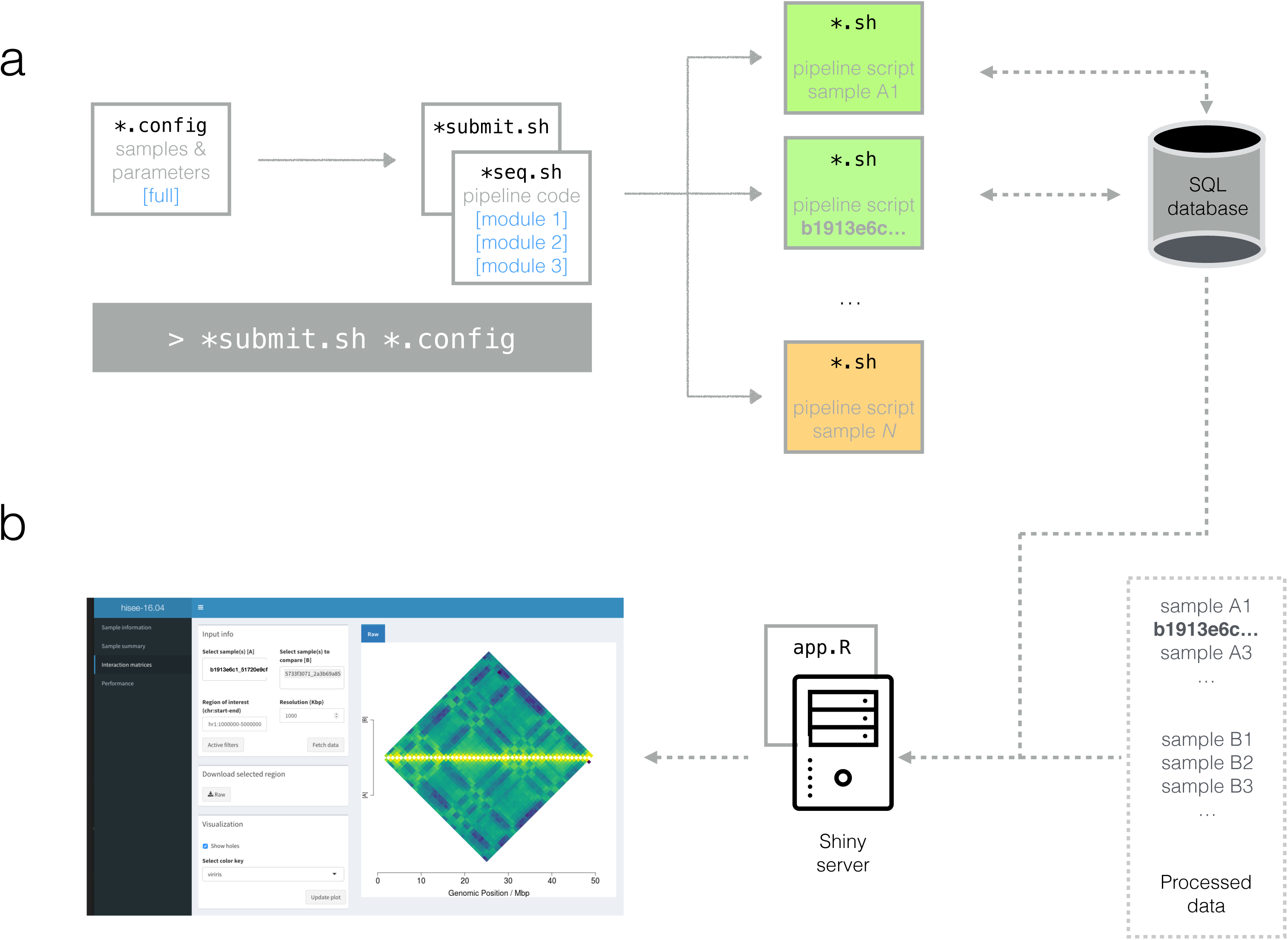
Automating the analysis and visualisation of b1913e6c1_51720e9cf data. **(a)** Scalability, parallelization, automatic configuration and modularity of analysis pipelines. Paul launched the Hi-C pipeline for hundreds of samples with a single command (gray rectangle): the submission script (“*.submit.sh”) generated as many pipeline scripts as samples listed in the configuration file (“*.config”). The configuration file also contained the hard-coded parameters shared by all samples, such as the maximum running time Paul underestimated for some samples. Processing hundreds of samples was relatively fast because (i) the pipeline script for each of the samples was submitted as an independent job in the computing cluster, where it was queued (orange) and eventually executed in parallel (green), and (ii) the pipeline code in “*seq.sh” was adapted for running in multiple processors. For further automation, each process retrieved sample-specific information (e.g. species, read length) from the metadata SQL database; in addition, metrics generated by the pipeline (e.g. running time, number of aligned reads) were recorded into the database. Because the pipeline code was grouped into modules, Paul was able to easily re-run the “generate_matrix” module for those samples that failed in his first attempt. **(b)** Interactive web application to visualise Hi-C data. b1913e6c1_51720e9cf alone generated ∼70 files of plots and text when passed through the Hi-C pipeline. Inspecting them might have seemed a daunting task for Chloe: she did not feel comfortable navigating the cluster and lacked the skills to manipulate them anyway, and even if she did, examining so many files for dozens of samples seemed endless. Luckily for her, Paul had developed and interactive web application with R Shiny (**Table 2**) that allowed her to visualise data and metadata and perform specific analyses in a user-friendly manner.

## Abbreviations

3K RGP: 3,000 Rice Genomes Project
ENCODE: Encyclopedia of DNA Elements
HTS: high-throughput sequencing
ID: identifier; SRA: Short Read Archive
SQL: Structured Query Language
TCGA: The Cancer Genome Atlas

## Declarations

### Ethics approval and consent to participate

Not applicable.

### Consent for publication

Not applicable.

### Availability of data and material

The didactic dataset is available at https://github.com/4DGenome/parallel_sequencing_lives

### Author’s contributions

Conceptualization: JQ, GF; Data curation: JQ; Formal analysis: JQ; Funding acquisition: TG, MAM-R, MB, GF; Methodology: JQ, EV, FD, YC, RS; Software: JQ, EV, FS; Visualisation: JQ, EV; Writing - original draft: JQ, GF; Writing - review & editing: EV, FD, FS, YC, RS, TG, MAM-R, MB. All authors read and approved the final manuscript.

## Acknowledgements

We thank F. Javier Carmona and Corey T. Watson for advice on the manuscript.

## Additional files

**Additional file 1. (a)** More than reads. FASTQ files may be useless if not coupled with biological, technical and logistics information (metadata). Metadata are used at several stages of the high throughput sequencing data. In the initial processing, for instance, the human origin of b1913e6c1_51720e9cf was needed to determine hg38 as the reference genome sequence to which reads would be aligned, and the restriction enzyme “DpnII” applied in the Hi-C protocol was used in the mapping too. Other metadata were used for quality control (e.g. sequencing facility and/or date for detecting batch effects or rescuing swapped samples using the correct index) or in the downstream analysis (e.g. cell type, treatment). Furthermore, metadata is critical for data sharing and reproducibility. **(b)** Choosing a name. Long before b1913e6c1_51720e9cf was generated, a scheme to name Hi-C samples was envisioned. First, two sets of either biological or technical fields that unequivocally defined a sequencing sample were identified. Then, for a given sample the values of the biological fields treated as text are concatenated and computationally digested into a 9-mer, and the same procedure is applied to the technical fields. The two 9-mers are combined to form the sample identifier (ID), as happened for b1913e6c1_51720e9cf. Despite the apparent non-informativeness of this sample ID approach, it easily allows identifying biological replicates and samples generated in the same batch since they will share, respectively, the first and second 9-mer. While the specific fields used to generate the sample ID can vary, it is important that they unambiguously define a sequencing sample (otherwise duplicated identifiers can emerge) and that they are always combined in the same order to ensure reproducibility. Indeed, another advantage of this naming scheme is that the integrity of the metadata can be checked, as altered metadata values will lead to a different sample ID.

**Additional file 2. Rapid accumulation and diversity of high throughput sequencing (HTS) data.** The past decade has witnessed a tremendous increase in sequencing throughput and applications, causing uncontrolled accumulation of sequencing datasets. **(a)** For instance, the number of sequences deposited in the Sequence Read Archive (SRA) [2], a major repository for HTS data, has skyrocketed from ∼2 Terabases in 2009 to ∼9,000 Terabases (the size of approximately 3 million human genomes) at the beginning of 2017. Moreover, this is surely an underestimation of the actual amount given that only sequencing experiments eventually included in a publication are deposited. Although data-intensive projects like TCGA [3], 1000 Genomes Project [4], ENCODE [5] and 3K RGP [6] are top HTS data generators [7], such a boost in the number of existing sequences reflects a pervasive use of HTS. **(b)** As an example, while sequencing data for >90,000 studies have been submitted to the SRA, the top 10 and 100 contributors in terms of number of bases represent only a part of the archive (∼30% and ∼60% respectively). **(c)** Similarly, while ∼80% of SRA data derive from *Homo sapiens* and *Mus musculus*, the central organisms in large sequencing projects, the remaining 20% come from a diverse number of organisms (∼50,000). Data were obtained from [8] and processed as described in the didactic dataset.

**Additional file 3. Why T47D_rep2 and b1913e6c1_51720e9cf are not singletons.**

**Additional file 4. Number of SRA deposited bases grouped by instrument name.** Data were obtained from [8] and processed as described in the didactic dataset.

**Additional file 5. Number of SRA deposited bases grouped by the submitter.** For the top 25 contributors in terms of number of bases submitted, we searched for instances of multiple entries probably referring to the same submitter (e.g. ‘ncbi’ and ‘NCBI’). Data were obtained from [8] and processed as described in the didactic dataset.

